# Assortment of *Stauntonia hexaphylla* and *Cornus officinalis* protect against testosterone-induced benign prostatic hyperplasia through anti-inflammatory and anti-proliferative activity

**DOI:** 10.1101/819474

**Authors:** Shanika Karunasagara, Geum-Lan Hong, Da-Young Jung, Kyung-Hyun Kim, Eun-Jeong Koh, Kyoung-won Cho, Sung-Sun Park, Ju-Young Jung

## Abstract

Benign prostatic hyperplasia (BPH) is a progressive pathological condition associated with proliferation of prostatic tissues, prostate enlargement, and lower-urinary tract symptoms. However, the mechanism underlying the pathogenesis of BPH is not clear. The aim of this study was to investigate the protective effects of *Stauntonia hexaphylla* and *Cornus officinalis* (SC extract) on a testosterone propionate (TP)-induced BPH model. For in vitro experiments, a human prostate adenocarcinoma cell line was used to perform western blotting for androgen receptor (AR), prostate specific antigen (PSA), and 5α-reductase type 2. Male Sprague-Dawley rats were randomly divided into 8 groups as follows for the in vivo experiments: control, BPH, Fina, Saw, SC25, SC50, SC100, and SC200. To induce BPH, all rats, except those in the control group, were daily administered with subcutaneous injections of TP (5 mg/kg), and orally treated with appropriate PBS/drugs for 4 consecutive weeks. Our findings indicated that the SC treatment significantly reduced the prostate size and downregulated the serum testosterone and DHT levels in BPH rats. The histological examination revealed that SC treatment markedly recovered the TP-induced abnormalities and reduced the prostatic hyperplasia. In addition, in vitro and in vivo western blotting indicated that SC treatment significantly downregulated the AR, PSA, and 5α-reductase type 2 expression, while an immunohistochemistry examination revealed that the SC extract significantly reduced the expression of type 2 5α-reductase and proliferating cell nuclear antigen positive cell count. Collectively, our findings demonstrated that SC extract attenuates BPH through anti-proliferative and anti-inflammation activities and might be useful in the clinical treatment of BPH.

## Introduction

Many plants have been identified as good sources of natural antioxidants, which protect against BPH and prostate cancer [1, 2]. In this study, we investigated the protective effect of a 9:1 mixture of *Stauntonia hexaphylla* and *Cornus officinalis*, called the SC extract, on BPH. *Stauntonia hexaphylla* belongs to the family Lardizabalaceae, which is native to Southern Japan and Korea and has been used in medicine owing to its analgesic, sedative, diuretic, and anti-cancer properties [3]. *Cornus officinalis*, native to Korea, Japan, and China comprises, compounds including terpenoids, flavonoids, sterols, carboxylic acids, polysaccharides, and phenylpropanoids that have been isolated and identified. Owing to its chemical constituents, *C. officinalis* displays diverse pharmacological activities such as hypoglycemic activity and protective activity toward diabetic target organs, and antioxidant, anti-inflammatory, and anticancer activity [4, 5].

Benign prostatic hyperplasia (BPH) is the most frequent, non-cutaneous form of cancer among elderly men and is characterized by progressive glandular and stromal tissue hyperplasia, which leads to an enlarged prostate [6]. The rapid growth of stromal and epithelial elements result in BPH in the prostate, along with lower urinary tract symptoms (LUTS), including obstructive symptoms such as hesitancy, poor intermittent stream, feeling of incomplete bladder emptying, and irritative symptoms such as frequency, urgency, and nocturia [7, 8]. Various molecular etiologies of BPH have suggested that hormones, oxidative stress, chronic inflammation, and aging may play a crucial role in BPH development, while researchers consider androgens, especially testosterone-related hormones to be the major contributory factors in the development and progression of BPH [9, 10]. The growth and development of prostate gland depends on androgen stimulation, especially through DHT [11, 12]; accumulation of DHT with aging results in rapid growth and hyperplasia of prostatic cells [13]. DHT is an active metabolic product of the conversion of testosterone by steroid 5α-reductase [14]. Finasteride, a drug that is used as a steroid 5α-reductase Type 2 inhibitor, can be used for the treatment for BPH [15]. It inhibits the conversion of testosterone to DHT, thereby preventing prostatic hyperplasia. Furthermore, saw palmetto has been widely used as a therapeutic remedy for urinary dysfunction due to BPH and works by ceasing the breakdown of testosterone into its byproduct DHT [16]. Race, family history of prostate cancer, and environmental factors act as possible risk factors of BPH [17-19].

The failure and unpleasant adverse effects of conventional anti-BPH drugs have led to the search for phytotherapeutic solutions as a safer and less toxic alternative. Recently phytotherapeutics have become popular in the treatment of BPH worldwide [20, 21]. This study was aimed to investigate the protective role of SC extract in the development of BPH in testosterone-induced BPH.

## Materials and method

### Plant material

SC extract (CKDHC-P29) was provided from Chong Kun Dang Healthcare (Seoul, Korea). Reference compounds, hederacoside D (purity ≥98.0%, ChemFace, Wuhan, China) and morroniside (purity ≥98.0%, ChemFace, Wuhan, China) were analyzed in the CKDHC-P29 sample. Reagents including acetonitrile, methanol, and ethanol were purchased from Burdick & Jackson (Muskegon, MI, USA) and formic acid (HPLC grade) was purchased from Sigma-Aldrich (St. Louis, Mo, USA). Water was purified using a Milli-Q system (Sinhan, Seoul, Korea).

### Preparation of SC extract

The SC extract consist of *Stauntonia hexaphylla* leaves (Lot number: 20181116) and *Cornus officinalis* Siebold & Zucc furits (Lot number: 20181112) mixed into 9:1 ratio. The leaves of *Stauntonia hexaphylla* were harvested in the area of Goheung-gun, Jeollanam-do, Korea and dried at 60 °C for 12h. Next, Dried leaves were extracted with 70% ethanol at 75°C for 12h and proceeded spray drying with 30% dextrin. The dried fruit of *Cornus officinalis* Siebold & Zucc fruit was harvested in the area of Gurye-gun, Jeollanam-do, Korea. Dried fruit was extracted with 70% ethanol at 75°C for 12h and proceeded spray drying with 50% dextrin.

### High performance liquid chromatography (HPLC) of SC extract

HPLC was used to identify the compounds, hederacoside D (C_53_H_86_O_22)_ and morroniside (C_17_H_25_O_11_) in the SC extract. The constituent of SC extract was performed by HPLC equipped with photodiode array detector (PDA, Berlin, Germany) using Agilent Zorbax eclipse plus C18 column (4.6 x 250 mm, 5 μm). The elution was performed using a linear gradient from 12 to 0.1% formic acid in acetonitrile for detection of hederacoside D and morroniside and the injection volume was 10 μL. PDA detector was set at 205 nm and 240 nm for appropriate hederacoside D (205 nm) and morroniside (240 nm).

### Cell culture and cell viability assay

Human prostate adenocarcinoma cells (LNCaP) were purchased from the American Type Culture Collection (ATCC, Manassas, VA, USA), seeded on to 6-well plates (5×105 cells/well) and fed with a medium containing 1 µM testosterone (Tokyo Chemical Ins. Co., Tokyo, Japan), and Finasteride (10 µM, Sigma, USA), Saw palmetto extract (100 µg/mL) or SC extract (25, 50 µg/mL). After 72 hours post treatment (72 hpt), cells were harvested for the preparation of total protein and subjected to western blot analysis. Cytotoxicity of SC extract on LNCaP cells was evaluated by MTT assay, using EZ-Cytox Cell Viability assay kit (Daeil Lab Service, Seoul, Korea) according to the manufacturer’s instruction. Cells were seeded at 1×104 cells per well and treated with concentrations of SC (0, 5, 10, 25, 50, 100 µg/mL) for 24 h. Absorbance was evaluated using microplate reader (BIO-TEK, Senergy HT), and cell viability was calculated as: 100% × (OD_450nm_ of SC group/ OD_450nm_ of control group).

### Experimental animals

The experimental protocols were approved (CNU-01108) by the International Animal Ethics Committee at Chungnam National University. Six-wk-old, male Sprague Dawley (SD) rats were purchased from www.orientbio.com and acclimated in specific-pathogen-free (SPF) animal facility under controlled temperature, humidity and photoperiod (22 ± 2 °C, 55 ± 5%, and 12h light/dark cycle respectively) for 1 wk before the experiment. All animals were fed with standard chow and water *ad libitum.*

### Experimental design

A total 40 SD rats were randomly divided into 8 groups (n = 5 /group) and treated for 4 wk as follows: Control group (PBS), BPH group (testosterone propionate/TP 5 mg/kg, S.C.), Fina (Finasteride 10 mg/kg, P.O.), Saw (Saw palmetto extract 100 mg/kg, P.O.), SC25 (SC extract 25 mg/kg, P.O.), SC50 (SC extract 50 mg/kg, P.O.), SC100 (SC extract 100 mg/kg, P.O.) and SC200 (SC extract 200 mg/kg, P.O.). To induce BPH, all rats except for the control group were daily given subcutaneous (S.C.) injections of TP (5 mg/kg) for 4 consecutive wk. To minimize the animal from suffering and distress while injection, S.C. injection carried out into the loose skin on the back of the neck and varied the site of injection to reduce the local skin reactions. At the end of the experimental period, rats were fasted overnight and euthanized by CO_2_ asphyxiation for 5 min in euthanasia apparatus. Blood was drawn from their abdominal vein. The blood samples were centrifuged at 3000 rpm at 4 °C, for 15 min and the serum was stored at −80 °C until further analysis. Finally, dissected prostate glands were weighed and stored at - 80 ° C liquid nitrogen for further analysis. Body weight of rats was measured at the beginning and the end of the experimental period.

### Assessment of serum testosterone and DHT levels

Serum testosterone and DHT were determined using commercial enzyme-linked immunosorbent assay (ELISA) kit (ALPCO Diagnostics, NH, USA) according to the manufacturer’s protocol.

### Histological study

Paraffin embedded prostate tissues were cut into 5µm sections. After deparaffinization and dehydration, sections were subjected to hematoxylin and eosin (H&E) staining and examined using light microscope (Nikson ECLIPSE Ni-U, Tokyo, Japan) at × 400 magnification. Images were taken from 10 randomly selected fields, and epithelial thickness was measured using Image J software (Image J v46a; NIH, USA).

### Western blot analysis of prostatic hyperplasia-related protein

Collected cells and frozen prostate tissue samples were lysed with RIPA buffer, centrifuged 12,000 rpm for 20 min at 4 °C and the supernatant was used for the western blotting. Extracted proteins were separated on 8-10 % SDS-polyacrylamide gels and transferred onto PVDF membrane using a semi-dry transfer system (Bio-Rad, Hercules, CA, USA). Then, primary antibodies were added as follows: anti-AR (1:1000, Santa Cruz), anti-PSA (1:1000, Santa Cruz) and anti-β-actin (1:5000, Abcam, Cambridge, UK). Horseradish peroxidase-conjugated goat anti-rabbit (AbFrontier, Seoul, Korea) was used to detect each protein and visualized with the enhanced chemiluminescence detection (ECL) kit (Amersham Pharmacia Biotech, Buckinghamshire, UK) and quantified using Image Lab Software (Bio-Rad, Hercules, CA, USA).

### Immunohistochemical (IHC) staining

After deparaffinization and dehydration, sections were subjected to 3% quenched endogenous H_2_O_2_ (in methanol), and treated with 0.5% Triton X-100 solution for 30 min at room temperature (25 ° C). Then, non-specific binding sites were blocked with normal goat serum (diluted 1:10 in PBS), and incubated overnight with primary antibodies as follows: anti-5α-reductase 2 (1:100; Santa Cruz) and anti-PCNA (Abcam, MA, U.S.A) at 4 ° C, incubated with the secondary antibody goat anti-rabbit (1: 200, Ab Frontier), and developed using diaminobenzidine (DAB) peroxidase substrate kit (Vector Laboratories). Stained sections were examined using light microscope (Nikon eclipse 80i, Nikon Corporation, Tokyo, Japan) at × 400 magnification. Images were captured by camera (Leica DCF450-C) at 10 different places of tissues, and 5-α reductase type 2, proliferating cell nuclear antigen (PCNA)-positive nuclei were counted.

### Statistical analysis

All data were expressed as mean ± standard deviation (SD). Statistical analysis was performed with t-test and Microsoft Excel 2016 was used as the statistical software. *p*-values less than 0.05 were considered statistically significant.

## Results

### HPLC chromatogram

Phytochemical compounds in the SC extract such as hederacoside D (Fig 1A) and morroniside (Fig 1C) were identified and quantified by using HPLC. Hederacoside D showed the characteristic peak at 39.52 min (Fig 1B), while morroniside at 6.53 min (Fig 1D) and the concentrations were found as 25 mg/g and 1.5 mg/g respectively.

**Fig 1.**
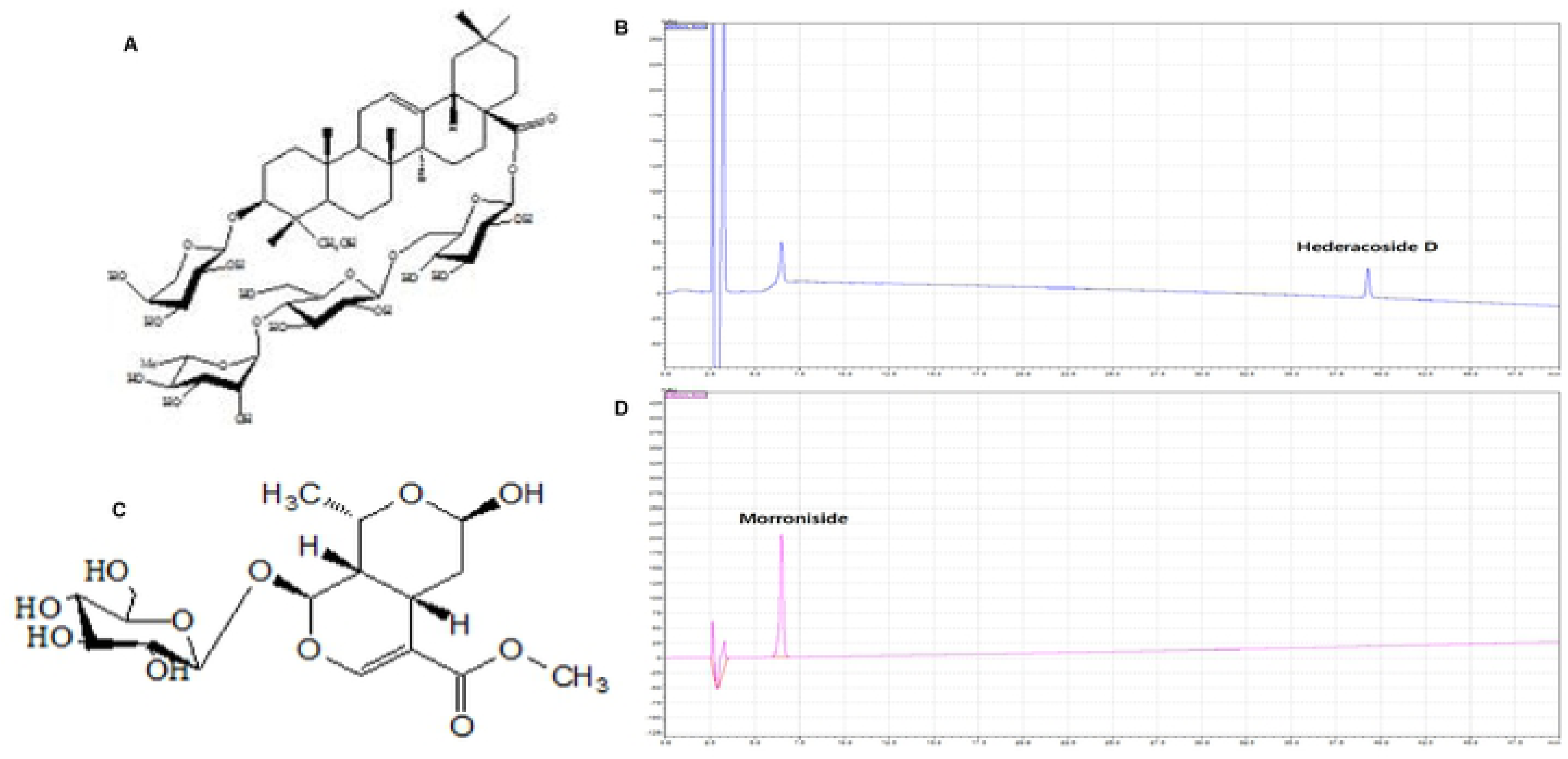
HPLC chromatogram of hederacoside D and morroniside. (A) Molecular structure of hederacoside D. (B) Constituent of hederacoside D in the SC extract was analyzed by HPLC-PDA. (C) Molecular structure of morroniside (D) Constituent of monroniside in the SC extract was analyzed by HPLC-PDA.

### Cell viability and in vitro western blotting

To study the cytotoxicity of SC extract on LNCaP cells, the MTT assay was performed. As shown in Fig. 2A, there was no significant difference in cell viability between SC treated cells and control cells. However, the exposure to 100 µg/mL of SC markedly decreased the cell viability up to 86.73%, demonstrating that high concentration of SC inhibits cell viability by its toxicity. Highest cell viability was shown by the cells treated with 50 µg/mL SC.

**Fig 2.**
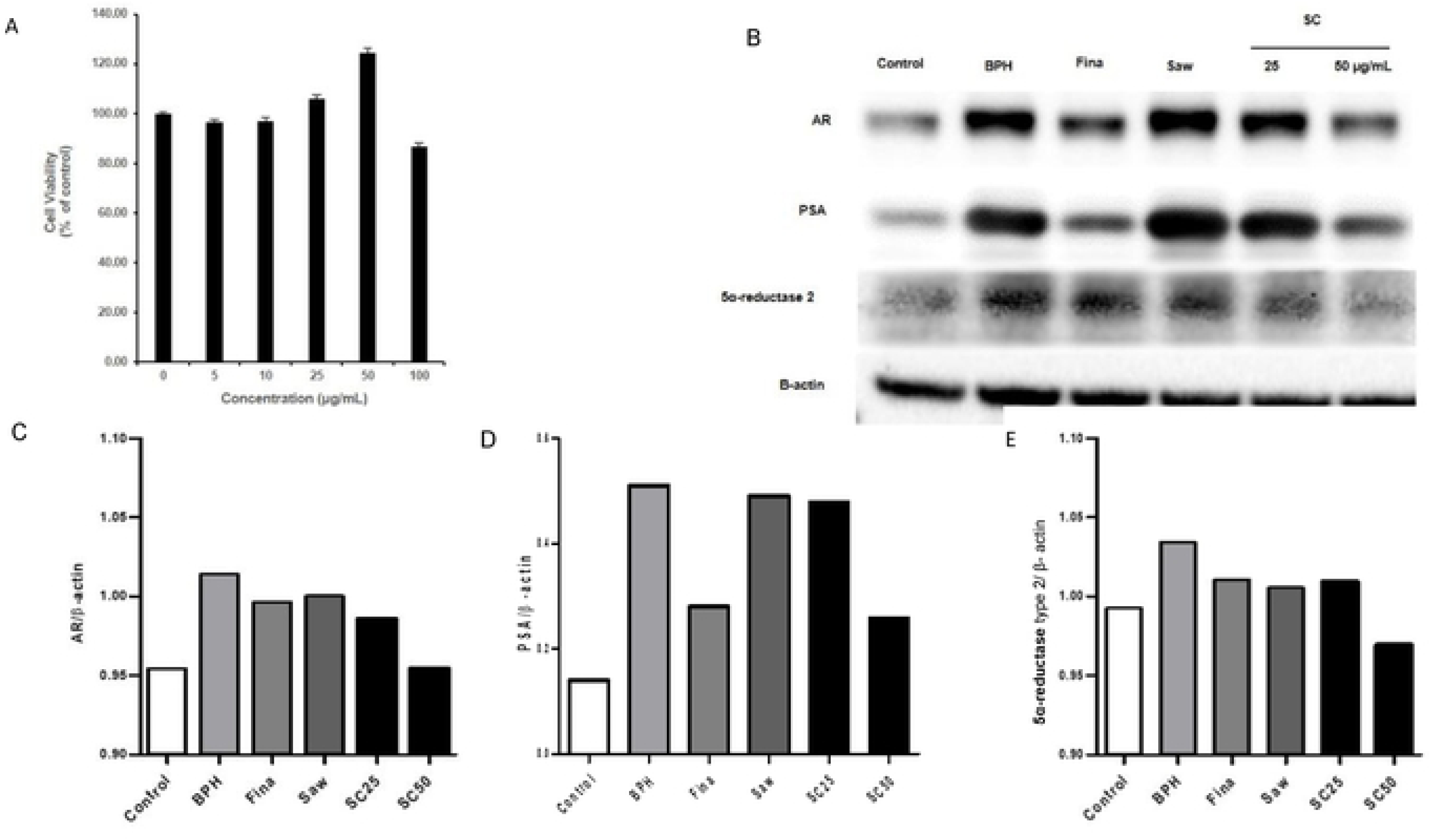
Effects of SC on the LNCaP human cancer cell line. (A) Cell viability LNCaP cells were treated with SC (0-100 μg/mL) for 72h, and cell viability was determined by MTT assay. (B) Western blot analysis of AR, PSA and 5α-reductasc type 2. LNCaP cells were incubated in medium containing testosterone (1 μM), Finasteride (10 μM), Saw palmetto (100 μg/mL) or SC (25,50 μg/mL) for 72h. (C, D, E) Relative densitometry evolution of AR, PSA and 5α-rcductasc type 2, the western blot data respectively.

In this study, we investigated the effect of SC treatment on androgen receptor (AR), prostate specific antigen (PSA), and 5α-reductase type 2 in LNCaP cells using western blot (Fig. 2B). As shown in Fig. 2C, D, E, TP treatment markedly increased the expression of AR, PSA and 5α-reductase type 2 in the BPH group relative to the control group. SC treated groups showed considerably decreased expression of aforementioned proteins, in a dose-dependent manner compared to the BPH group. Further, expression of AR, PSA and 5α-reductase type 2 in the finasteride and saw palmetto treated groups exhibited decreased expression compared to the BPH group. However, there was no significant different between the control and BPH-induced groups. These findings suggest that SC treatment suppresses androgen signaling in LNCaP cells.

### Effect of SC extract on body weight and prostate weight

Examination of body weight (BW) and prostate weight (PW) is commonly use to evaluate the progression of BPH. As shown in Fig. 3A, rats in the TP-induced BPH group showed significantly (*p* < 0.001) increased PW compared to the control group. The rats treated with finasteride showed significantly (*p* < 0.01) decreased PW relative to the BPH group. Similarly, SC50 and SC200 groups also exhibited significantly (*p* < 0.001, *p* < 0.01 respectively) decreased PW compared with the BPH group. However, no significant differences in BW were observed among the groups (Fig. 3B). Moreover, relative prostatic weight was significantly (*p* < 0.001) elevated in the BPH group compared to the control, while Fina group exhibited significantly (*p* < 0.001) decreased values in comparison to the BPH group (Fig. 3C). The groups SC50 and SC200 also showed these values that were significantly (*p* < 0.001, *p* < 0.01respectively) decreased than those of the rats in the BPH group. These results proved that SC is a potential treatment for modulating the prostate size of the TP-induced BPH model.

**Fig 3.**
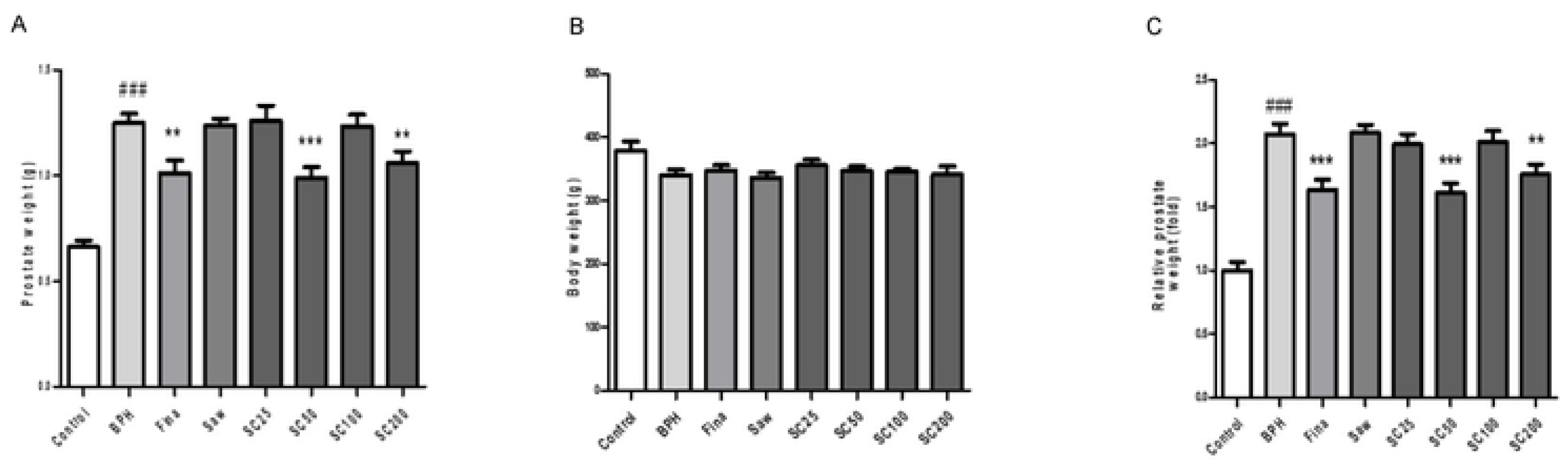
Effect of SC on prostate weight and body weight in TP-induced BPH rats. Abbreviations: Control (PBS), BPH (5 mg/kg TP, S.C.), Fina (Finasteride 10 mg/kg. P.O.), Saw (Saw palmetto 100 mg/kg, P.O.), SC25 (SC extract 25 mg/kg, P.O.), SC50 (SC extract 50 mg/kg. P.O.), SC100 (SC extract 100 mg/kg, P.O.) and SC200 (SC extract 200 mg/kg. P.O.) To induce BPH, all rats were received daily TP injection (5 mg/kg, S.C.) for consecutive 4 wk, except for the control group. (A) Prostate weight, (B) Body weight, (C) Relative prostate weight. Data are expressed as the means ± S.E.M (n=6). ###*p* < 0.001 versus control group, \*\**p* <0.01 versus BPH group, *** *p* < 0.001 versus BPH group.

### SC mediated downregulation of serum testosterone and DHT in BPH rats

In this experiment, we studied serum testosterone and DHT as they have a fundamental role in progression of BPH. As shown in Fig. 4A, a significant (*p* < 0.01) increase in serum testosterone level was shown in the BPH group compared with the control group. In contrast, rats in the groups of Fina, SC50 and SC200 showed significantly (*p* < 0.01) decreased testosterone levels relative to the BPH group. Similarly, the groups Saw, SC25 and SC100 also showed significant (*p* < 0.05) increase in the testosterone level. Like the serum testosterone, serum DHT level also significantly (*p* < 0.01) increased in the BPH group (Fig. 4B). By contrast, the rats in all the SC treated groups showed significantly (*p* < 0.001) reduced DHT level. Like rats in the SC treated groups, those in the finasteride treated group also showed significantly (*p* < 0.05) increased serum DHT level. These data evidenced that SC extract reduced the androgen concentrations in BPH induced rats.

**Fig 4.**
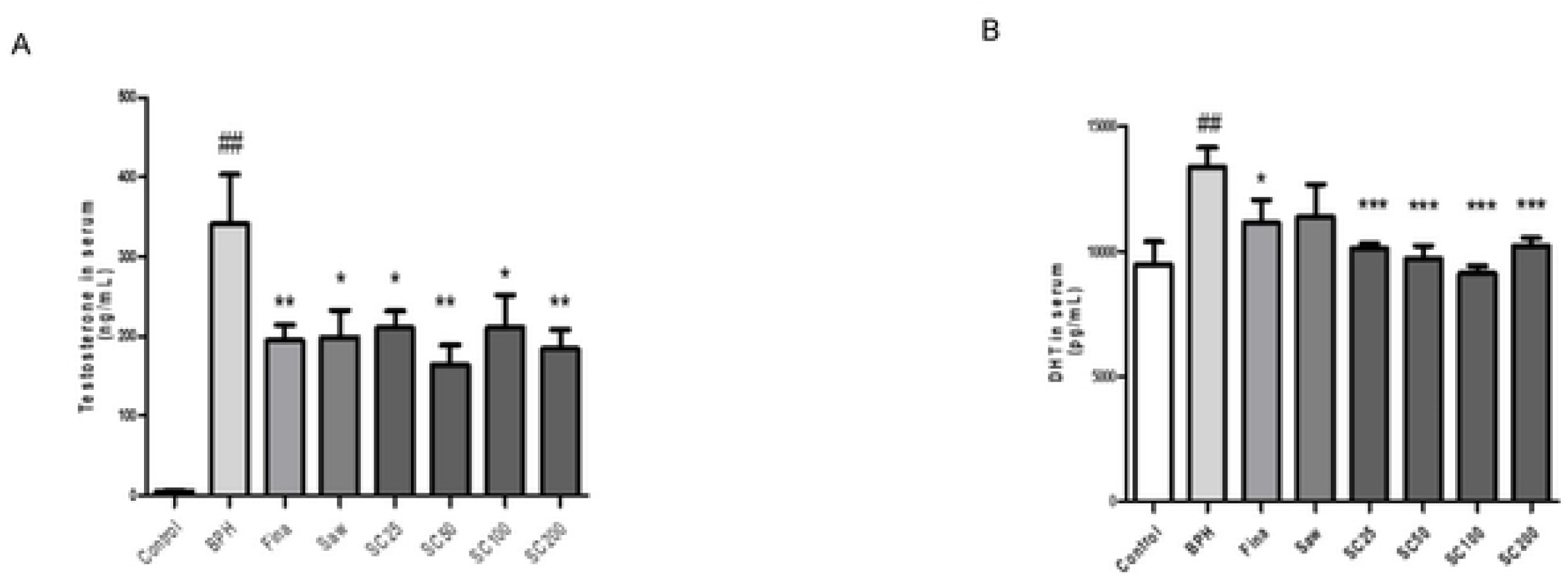
Effect of SC on scrum testosterone and DHT in TP-induccd BPH rats. Individual data were obtained using an ELISA assay. Abbreviations Control (PBS). BPH (5 mg/kg TP, S.C.), Fina (Finasteride 10 mg/kg, P.O.), Saw (Saw palmetto 100 mg/kg, P.O.), SC25 (SC extract 25 mg/kg, P.O.), SC50 (SC extract 50 mg/kg, P.O.), SC100 (SC extract 100 mg/kg, P.O.) and SC200 (SC extract 200 mg/kg, P.O.) To induce BPH, all rats were received daily TP injection (5 mg/kg, S.C.) for consecutive 4 wk, except for the control group. (A) The serum concentrations of testosterone. (B) The serum concentrations of DHT Data are expressed as the means ± S.E.M (n=6). ##*p* < 0.01 versus control group. \**p* <0.05 versus BPH group. \*\**p* <0.01 versus BPH group. \*\*\**p*< 0 001 versus BPH group.

### Study of morphological changes of prostate tissue in BPH rats

To demonstrate the TP-induced histological alterations, H&E staining was carried out (Fig. 5A). There were no histological changes associated with the control group. However, BPH group showed disrupted morphology such as significant thickening, hypertrophy and hyperplasia with papillary projections in the epithelium. Further, the lumen diameter was also widened as compared to the control group. Mild epithelial hyperplasia was shown by both Fina and Saw groups relative to the BPH group. SC treatment has been shown to markedly reduce the above morphological abnormalities in the BPH rats, in a dose dependent manner. As shown in Fig, 5B, rats in the BPH group exhibited significantly (*p* < 0.001) increased prostatic epithelial thickness compared with those of the control group. In contrast, all the BPH rats, which have been treated, showed significant (*p* < 0.001) reduction in prostatic epithelial thickness compared to the control group. Aforementioned results suggest that SC treatment has an ability to attenuate the abnormal histological changes, thereby reduce the prostate weight in the BPH rats.

**Fig 5.**
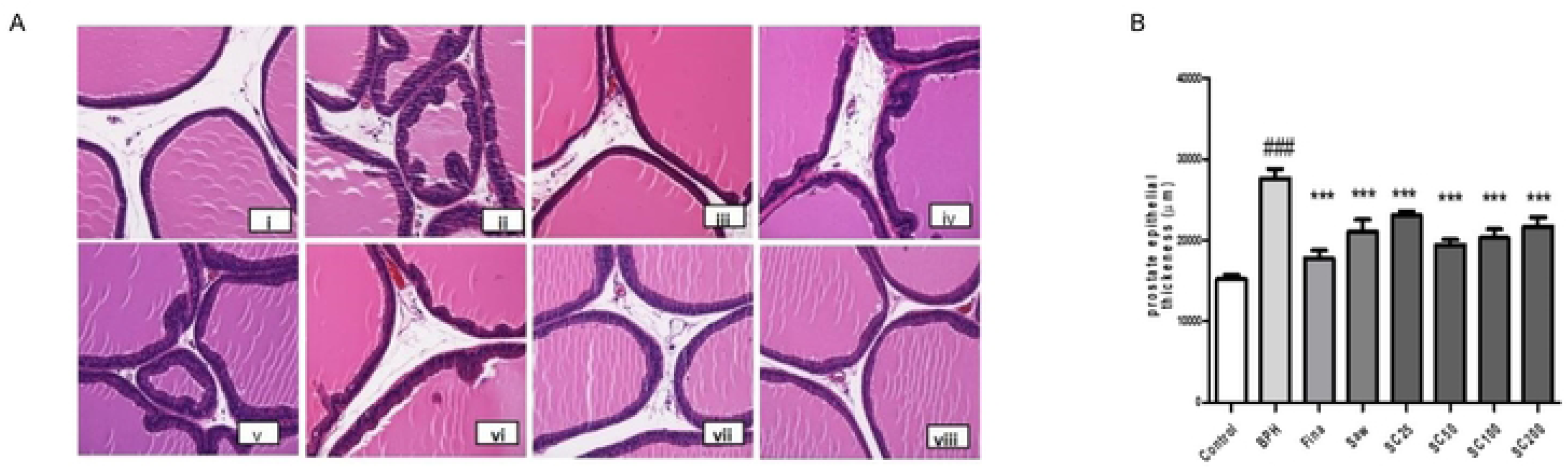
Effects of SC on histological changes in prostate tissues in TP-induced BPH rats. Prostate tissues were stained with H&E solution for histological study (magnification, 400×). Abbreviations Control (PBS). BPH (5 mg/kg TP. S.C.), Fina (Finasteride 10 mg/kg. P.O.), Saw (Saw palmetto 100 mg/kg. P.O.), SC25 (SC extract 25 mg/kg, P.O.), SC50(SC extract 50 mg/kg, P.O.), SC100 (SC extract 100 mg/kg. P.O.) and SC200 (SC extract 200 mg/kg, P.O.) To induce BPH, all rats were received daily TP injection (5 mg/kg, S.C.) for consecutive 4 wk, except for the control group. (A) Representative photomicrographs of prostate sections. (B) Prostate epithelial thickness. (i) Control; (ii) BPH; (iii) Fina; (iv) Saw; (v) SC25; (vi) SC50; (vii) SC100; (SC200). ###*p*< 0.001 versus control group, \*\*\**p*< 0.001 versus BPH group.

### In vivo western blotting

For in vitro analysis, western blotting was performed to evaluate AR and PSA protein expression in the prostate tissue (Fig. 6A). As shown in Fig. 6 B, C, AR exhibited a significant (*p* < 0.01) increase and PSA showed markedly increased expression in the BPH group compared with the control group. The rats treated with finasteride and SC, showed significantly (*p* < 0.05) reduced expression of AR and markedly decreased PSA expression relative to the BPH group. These results indicated that SC treatment has an ability to suppress the androgen signaling in prostate cells of BPH rats.

**Fig 6.**
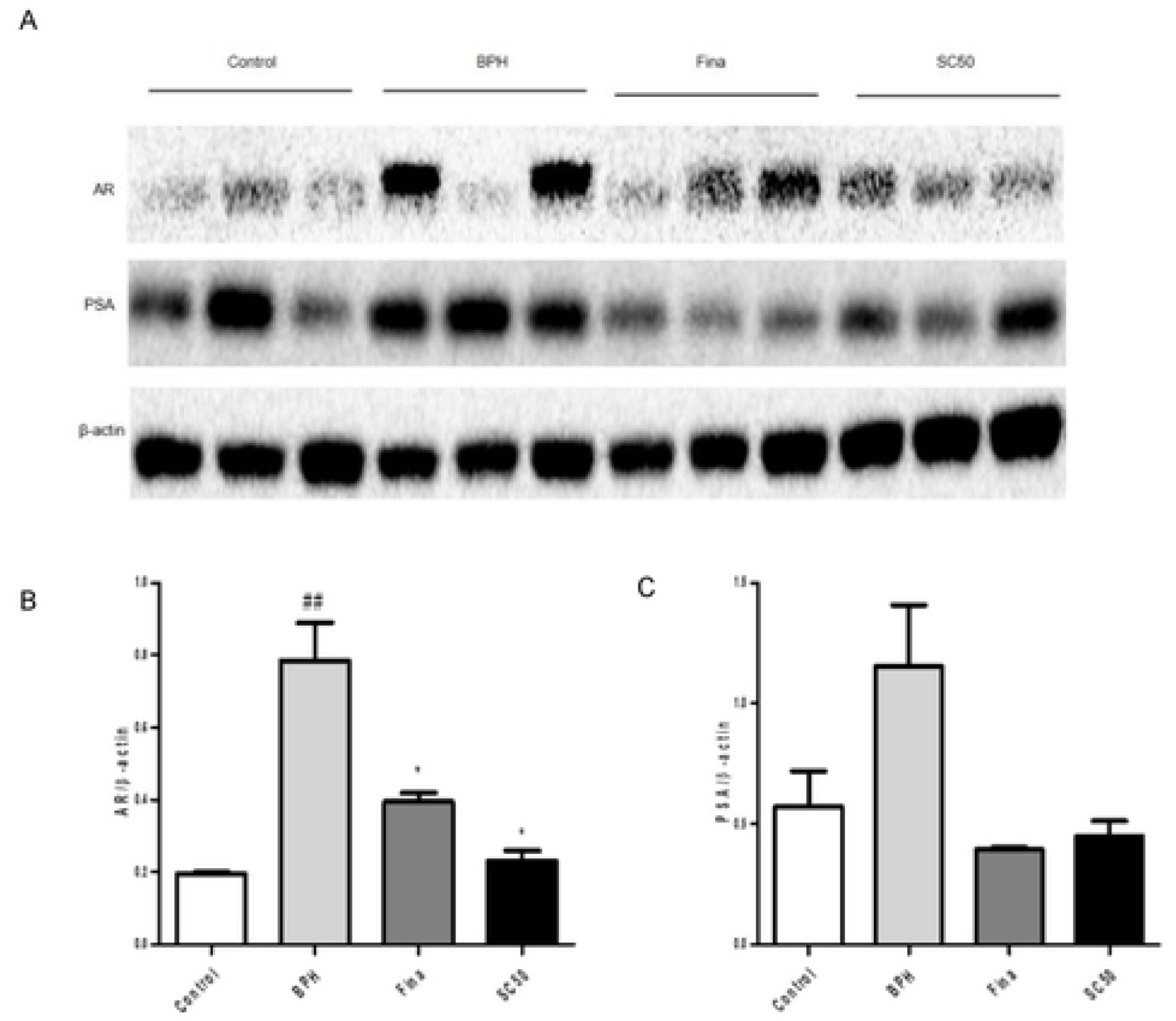
Effect of SC on prostatic AR and PSA in TP-induced BPH rats. Abbreviations: Control (PBS), BPH (5 mg/kg TP, S.C), Fina (Finasteride 10 mg/kg. P.O.), Saw (Saw palmetto 100 mg/kg, P.O.), SC25 (SC extract 25 mg/kg, P.O.), SC50 (SC extract 50 mg/kg, P.O.), SC100 (SC extract 100 mg/kg, P.O.) and SC200 (SC extract 200 mg/kg, P.O.) To induce BPH. all rats were received daily TP injection (5 mg/kg, S.C.) for consecutive 4 wk, except for the control group. (A) Western blot analysis of prostatic AR and PSA. (B, C) Relative densitometry evolution of AR and PSA, the western blot data.

### Immunohistochemical study of rat prostatic tissue

In this study, we investigated the immunoexpression of 5α-reductase type 2 and PCNA. As shown in Fig. 7A, IHC of prostate tissue indicated that the rats in the control group had no abnormalities; however, those in the BPH and Saw groups showed increased expression of type-2 5α-reductase. Notably, finasteride and SC treatment reversed the effect of TP, and SC treatment reduced the type-2 5α-reductase expression, in a dose-dependent manner. We also studied the anti-proliferative effect of SC by measuring the expression of PCNA in the prostate tissues and PCNA-positive cells were characterized by brown-stained nuclei, in both glandular epithelium and stroma. As shown in Fig. 7B, rats in the control group had no expression of prostatic epithelial PCNA, however rats in the BPH and Saw groups exhibited an increased number of PCNA-positive cells. Finasteride and SC partially reversed the effect of TP. Moreover, as shown in Fig. 7C, BPH group exhibited significantly (*p* < 0.05) increased PCNA expression relative to the control group, while finasteride and SC treatments significantly (*p* < 0.05, *p* < 0.01 respectively) reversed the effect of TP as compared to the BPH group. These data proved that SC extract prevents BPH through anti-proliferative activity.

**Fig 7.**
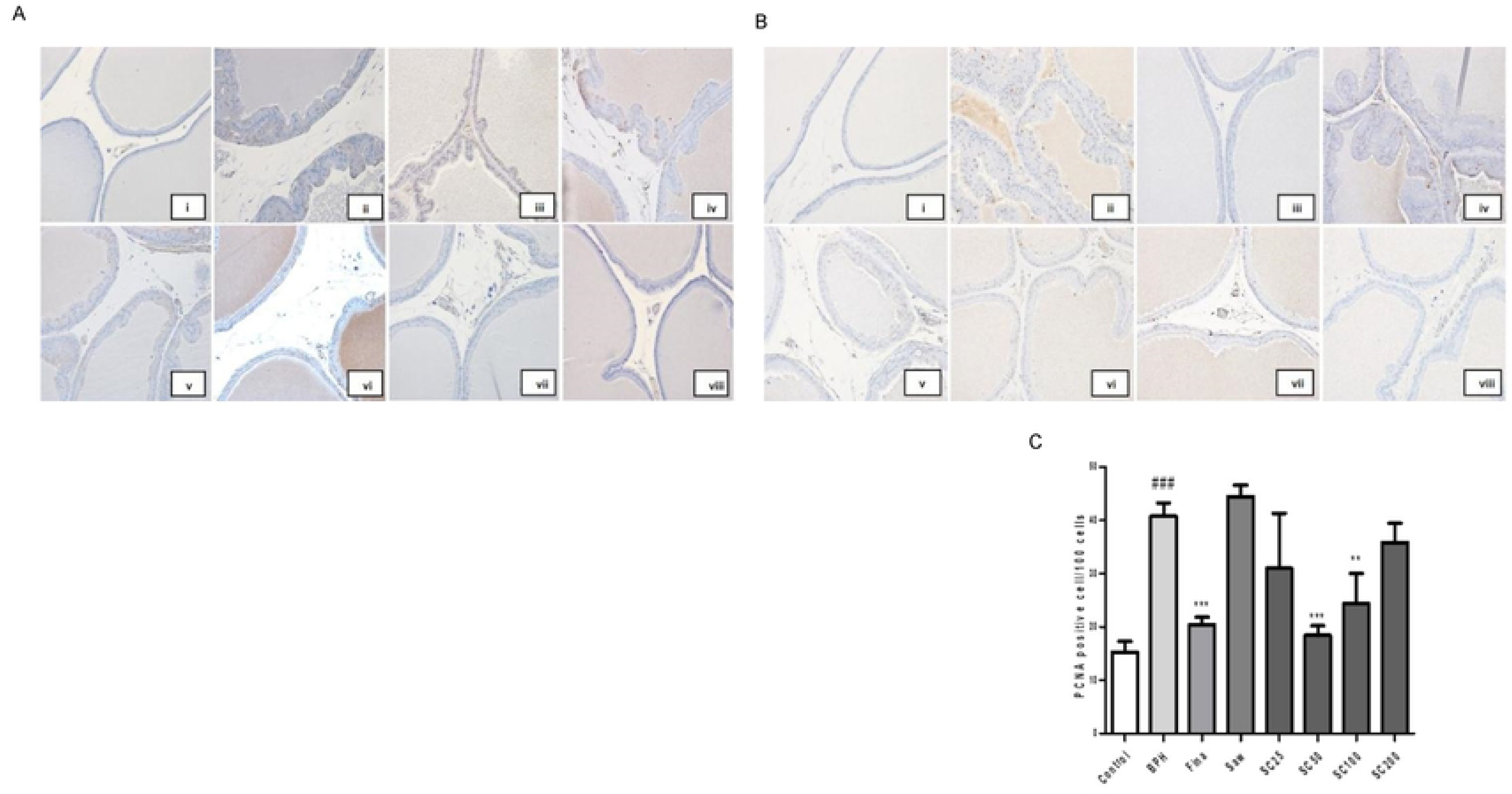
Effect of SC on type 2 5α-reductase and prostate cell proliferation in TP-induced BPH rats. Prostate tissues were immune stained with anti-5α-reductase and anti-PCNA antibody. PCNA positive nuclei were counted in 10 randomly selected fields (magnification. 400×). Abbreviations Control (PBS), BPH (5 mg/kg TP. S.C), Fina (Finasteride 10 mg/kg, P.O.). Saw (Saw palmetto 100 mg/kg, P.O.), SC25 (SC extract 25 mg/kg, P.O.), SC50 (SC extract 50 mg/kg, P.O.), SC100 (SC extract 100 mg/kg, P.O.) and SC200 (SC extract 200 mg/kg, P.O.) To induce BPH, all rats were received daily TP injection (5 mg/kg, S.C.) for consecutive 4 wk, except for the control group. (A) Representative photomicrographs of type 2 5α-rcductase expression. (B) Representative photomicrographs of PCNA expression. (C) PCNA positive nuclei/100 cells. (i) Control; (ii) BPH; (iii) Fina; (iv) Saw; (v) SC25; (vi) SC50; (vii) SCI00; (SC200). ###*p* < 0.001 versus control group, \*\**p* <0.01 versus BPH group,\*\*\**p*< 0.001 versus BPH group

## Discussion

Benign prostatic hyperplasia and associated lower urinary tract symptoms are common urological problems among older men, and sex steroids have a fundamental role in the development and maintain of BPH. This study may provide an alternative therapeutic option to BPH by using herbals alternatives. We found that SC treatment could significantly attenuate the development and progression of TP-induced BPH, which was confirmed by decreased PW, relative prostate weight and histological changes. This was further proved by western blot and IHC data.

BPH is a noncancerous prostate condition, which is caused by the overgrowth of prostatic epithelial and stromal cells, and arises due to the imbalance between proliferation and apoptosis of prostatic cells [22]. Increased proliferation of epithelial and stromal cells in the prostate cause the overall enlargement of the prostate gland, which is a signal of BPH development [23, 24]. Hyperplasia of these cells results in increased prostate weight, which causes contraction of the urethral canal, which may obstruct the urine flow [25]. In line with previous studies [23, 26], our investigation also demonstrated that rats with experimentally induced BPH have significantly higher PW relative to the control group, while finasteride and SC treatments have significantly inhibited the prostate weight gain as compared to the BPH group. Our histopathological examination demonstrated that BPH group had a significant epithelial hyperplasia and epithelial thickness relative to the control, while finasteride, saw palmetto and SC treatments significantly inhibited the above condition as compared to the group with no treatments. These results indicated that SC could be an alternative treatment for prostatic hyperplasia in TP-induced BPH.

Testosterone is an androgen produced by Leydig cells of the testis, while DHT is an important byproduct of testosterone and the most potent androgen in men, which is involved in development and aging. Further, type 2 5α-reductase is considered to be as the convertor of testosterone to its metabolite DHT [27-31]. Our study demonstrated significantly increased testosterone and DHT levels in the serum of BPH group as compared to the control. On the other hand, a significant reduction was observed in aforementioned androgens in the finasteride and SC treated rats as compared to the BPH group. These data evidenced that SC treatment has an ability to regulate androgens and their byproducts, and thereby attenuate BPH development.

Androgens affect gene expression in various kinds of tissues and cells by binding with the AR, which has been linked to prostate cancer [32, 33]. DHT has a higher affinity for AR as compared to testosterone; and in the prostate, the interaction between DHT and AR results in the production of proteins such as PSA. PSA is a glycoprotein in humans, encoded by the KLK3 gene, a member of the kallikrein-related peptidase family, and secreted by the prostatic epithelial cells. It plays a role in various functions during copulation and fertilization [34]. As serum PSA levels are frequently raised in prostate disorders such as BPH and prostate cancer, it is used as a clinical maker for disease prognosis [35]. In this study, in vitro and in vivo western blotting revealed that the increased AR, PSA and 5α-reductase type 2 in BPH condition can be significantly inhibited with finasteride and SC treatments. These findings proved that SC extract is a potential option to suppress androgen signaling in prostatic cells.

In the current study, we also performed the IHC to examine the expression of 5α-reductase type 2 and PCNA. PCNA is a marker of cell proliferation, particularly expressed in proliferating cell nuclei, and plays a crucial role in certain physiological and pathological conditions. Disruption of balance between cell death and cell proliferation can cause abnormal growth of prostate gland, leading to BPH [36, 37]. Oxidative stress is also mediated by the mechanisms that are associated with prostate proliferation [38]. In this study, significantly increased PCNA expression was detected in the BPH group rats as compared to the control rats. In contrast, both finasteride and SC treated rats exhibited significant reduction in PCNA count relative to the BPH group. Moreover, IHC of type 2 5α-reductase showed elevated expression in BPH group, while both finasteride and SC intervention reversed that effect. Together, these findings proved that SC extract can be a possible treatment for BPH via anti-proliferative activity.

## Conclusions

In conclusion, the findings of our study revealed that SC treatment significantly reduced the prostate hyperplasia and prostate size. In addition, SC extract had an ability to inhibit type 2 5α-reductase expression, thereby preventing the conversion of testosterone into its byproduct DHT, suggesting that SC is a potential treatment for regulating androgen signaling in prostatic cells. Moreover, intervention of SC resulted in anti-proliferative activity, thereby prevent the development and progression of BPH. Therefore, this investigation evidenced that assortment of *Stauntonia hexaphylla* and *Cornus officinalis* can protect against testosterone-induced benign prostatic hyperplasia through its anti-inflammatory and anti-proliferative activity. Use of SC extract as a therapeutic intervention for the treatment of BPH warrants further investigation.

## Data availability

The data used to support the finding of this experiment are available from the corresponding author upon request.

## Acknowledgement

This research was supported by Korea Institute of Planning and Evaluation for Technology in Food, Agriculture and Forestry (IPET) through High Value-added Food Technology Development Program.

## Author contribution

**Conceptualization:** Ju-Young Jung; **Data curation:** Geum-Lan Hong; **Funding acquisition:** Ju-Young Jung; **Investigation:** Shanika Karunasagara, Geum-Lan Hong, Da-Young Jung, Eun-Jeong Koh, Kyoung-won Cho, Sung-Sun Park; **Supervision:** Ju-Young Jung; **Validation:** Da-Young Jung, **Writing-original draft:** Shanika Karunasagara, **Writing-review & editing:** Ju-Young Jung

## Abbreviations

BPH: Benign prostatic hyperplasia;
LUST: lower urinary tract symptoms;
TP: testosterone propionate,
DHT: dihydrotestosterone;
AR: androgen receptor;
PSA: prostate specific antigen;
PCNA: proliferating cell nuclear antigen;
S.C.: subcutaneous;
P.O.: Per os/ oral administration.

